# *biomes*: An R package for reproducible occurrence-to-biome classification using 31 global biome schemes

**DOI:** 10.64898/2026.09.25.754345

**Authors:** Hans Christian Groß, Jan-Christopher Fischer, Anna Walentowitz, Alexander Zizka

**Affiliations:** Department of Biology, University of Marburg, Karl-von-Frisch-Straße 8, 35043 Marburg, Germany; Department of Biogeography, University of Bayreuth, Universitätsstraße 30, 95447 Bayreuth, Germany; Bayreuth Centre for Ecology and Environmental Research (BayCEER), Universitätsstraße 30, 95447 Bayreuth, Germany; Marburg Center for Biodiversity Monitoring and Modelling (M3B), Karl-von-Frisch-Straße 8, 35043 Marburg

**Keywords:** biogeography, biome scheme, GBIF, macroecology, occurrence record, reproducibility, vegetation classification

## Abstract

Biomes are continental-scale vegetation units widely used in ecological and evolutionary research, often together with large-scale species-distribution data. Many conceptually distinct biome schemes exist, differing in the underlying definition, delineation methodology and spatial resolution, yet most studies using biomes default to a few readily available schemes rather than choosing the one best suited to the research question.
We present *biomes*, an R package distributing spatially explicit raster layers of 31 published global biome schemes in a harmonised format at a 10 x 10 km resolution, together with metadata and a concise description. To facilitate their use, *biomes* provides a user-friendly classification workflow to classify occurrence records into specific biomes and a data-driven ranking algorithm to help researchers to choose the most suitable biome scheme for their data and research question in a transparent, reproducible way.
The *biomes* workflow can be applied to any dataset of georeferenced terrestrial occurrences. It (1) accepts either user-provided occurrence records or records downloaded by the package, (2) ranks the schemes according to their fit to the occurrence data, (3) assigns the records to specific biomes, and (4) summarises the resulting taxon-biome patterns. We demonstrate this workflow using 17,030 occurrence records of 185 species of Bombacoideae (Malvaceae). The analysis identified the vegetation-based biome scheme of Ramankutty and Foley (1999) as the best fit and recovered the forest and savanna biome signal known in the group from the literature, with tropical evergreen woodland being the biome with the largest share of records (46.1%) and highest number of species (162 of 185 species; 87.6%).
*biomes* provides user-friendly access to technically standardised and spatially explicit layers of 31 biome schemes, and thereby enables the transparent, data-driven choice of biome schemes tailored to a specific research data and questions, improving the reproducibility and the specificity of biome use across ecological and evolutionary research.

**Data and code for peer review:** the development version of the *biomes* package is available at a GitHub (https://github.com/azizka/biomes); the stable release is available from CRAN (https://cran.r-project.org/web/packages/biomes).

## Introduction

Biomes are continent-level vegetation units, characterised by vegetation physiognomy and broad-scale environmental conditions, and are widely used to partition the terrestrial biosphere in research and education (Fischer et al., 2022; Mucina, 2019). However, three practical challenges complicate the use of biomes in quantitative research. First, although Fischer et al. (2022) consolidated the underlying digital layers into a harmonised raster stack, the stack remains a standalone data product and is not available through a dedicated R package. Given the widespread use of R in ecological research, package-based access would lower the technical barriers to using these data and allow researchers to retrieve and process them directly within reproducible analytical workflows. Second, there is no transparent way to choose among competing biome schemes, even though this choice can affect downstream inference (Scheiter et al., 2024). Third, there is no reproducible way to assign species occurrence records to individual biomes within a selected biome scheme. Instead, such classification is typically performed ad hoc using bespoke, potentially error-prone code, with inconsistent treatment of records that fall outside the scope of the selected biome scheme. In practice, these limitations lead most studies to default to one of a few readily available biome schemes rather than the one best suited to the specific research question addressed. The *biomes* package solves these issues by providing georeferenced GIS layers for 31 biome schemes at a 10 x 10 km resolution in a global, harmonised format, which makes them particularly suited to macroecological and large-scale biogeographical analyses.

The choice of biome scheme is crucial for most analyses. Biome schemes differ conceptually in how they define a biome and, as a result, also in the resulting spatial delineation of specific biomes. Biomes may conceptually be defined by climate (resulting for instance in specific biomes such as “polar frost”), by physiognomy (“savanna”), by vegetation functional traits (“evergreen forest”), or by combinations thereof (“moist evergreen broadleaf rainforest”). As a result, a plethora of schemes based on different definitions has been proposed, ranging from classical climate envelopes (Schimper, 1898; Whittaker, 1970) through assemblies of ecoregions based on expert-judgment (Dinerstein et al., 2017; Olson et al., 2001) and model-derived vegetation maps (Allen et al., 2020; Conradi et al., 2020) to remote-sensing land-cover products (Friedl et al., 2010). Following their definition, these schemes differ in the underlying data, methodology of biome delineation, spatial resolution and extent. Therefore, using an ill-suited scheme can distort inferences from research outputs in several ways. First, there is a risk of circular reasoning when the variable(s) used to define biomes, coincide(s) with the variable(s) under study in subsequent analysis. For instance, using a trait-based scheme to test the distribution of plant functional traits builds the expected signal into the classification. Second, the data underlying biome delineation vary in spatial coverage, resolution, and quality, resulting in geographically uneven levels of detail and uncertainty in biome maps. Consequently, biomes may unduly merge or fragment ecologically meaningful structures relative to a specific study system and research question. For instance, a coarse-resolution global scheme may assign an entire elevational gradient to a single forest biome, thereby obscuring physiognomic and environmental differences between lowland and montane forests. Such scale mismatches can bias downstream analyses by treating ecologically distinct areas as equivalent. Third, in the common application of assigning occurrences to biome, unsuitable resolution and spatial coverage can cause data loss. For instance, if the spatial resolution of a particular biome scheme is low, coastline records may be located outside the defined biomes and thus require time consuming manual curation. Hence, the choice of biome scheme will affect the inferences drawn and therefore, ideally, an appropriate biome scheme should be selected for any research question (Fischer et al., 2022; Scheiter et al., 2024). However, differences among global biome schemes can be difficult for users to recognise, and familiar classifications are therefore often adopted uncritically (Beierkuhnlein & Fischer, 2021).

The *biomes* package addresses these three challenges by (1) providing user-friendly access to 31 harmonised global biome schemes (Fischer et al., 2022) in a standardised format from within R; (2) providing guidance in choosing the most suitable biome scheme for a given dataset and research question through a data-driven ranking on three complementary criteria: spatial coverage, effective number of occupied biomes, and granularity; and (3) standardising occurrence-to-biome classification of geo-referenced occurrence records in a reproducible and user-friendly way.

### Installation and usage

The development version of biomes is available from GitHub (https://github.com/azizka/biomes) a stable release is available from CRAN (https://cran.r-project.org/web/packages/biomes/). Package structure, coding style, and documentation follow the conventions of the rOpenSci package development guide (rOpenSci et al., 2024). The package requires R (≥ 4.1.0; R Core Team, 2026) and depends on the packages *terra* (Hijmans et al., 2026), *readr* (Wickham, Hester, et al., 2026), *checkmate* (Lang et al., 2026), *sf* (Pebesma et al., 2026), *ggplot2* (Wickham, 2016; Wickham, Chang, et al., 2026), *viridis* (Garnier et al., 2024), *tidyterra* (Hernangómez, 2023; Hernangómez et al., 2026) and *rlang* (Henry & Wickham, 2026). Further packages suggested for the vignettes and for optional functionalities are: *dplyr* and *tidyr* (Wickham et al., 2025; Wickham, François, et al., 2026), *cowplot* (Wilke, 2025) and *ggforce* (Pedersen & RStudio, 2025). *rgbif* (occ_search() and occ_download(); Chamberlain et al., 2026; Chamberlain & Boettiger, 2017) and *CoordinateCleaner* (clean_coordinates(); Zizka et al., 2019).

### The ***biomes*** workflow

In practice, *biomes* provides a four-step-workflow (Fig. 1). Each step corresponds to one function of the package, and the high-level wrapper biomes_full() runs the entire pipeline in a single call for a taxon name. Below we provide a worked example using a global dataset comprising 17,030 occurrence records for 185 species of Bombacoideae (Malvaceae) as a worked example. Further documentation is available in the package documentation and the accompanying vignettes (https://azizka.github.io/biomes/index).

**Figure 1.**
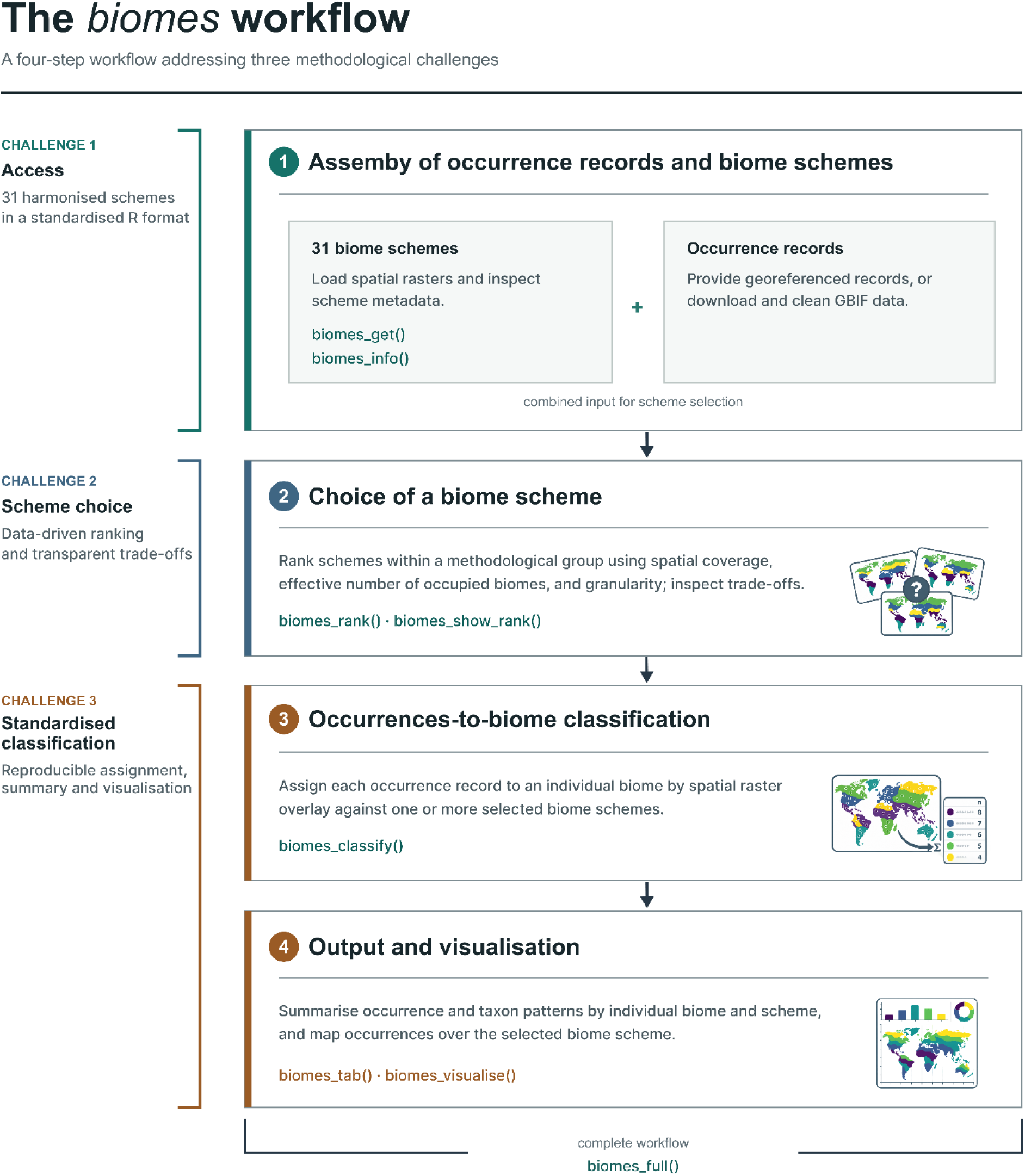
The standard *biomes* workflow with four steps. Each box describes one step including the respective package functions. Steps are mapped on the three challenges to the use of biomes in Ecology and evolution.

#### Step 1: Assemby of occurrence records and biome schemes

biomes_get() provides access to 31 biome schemes (Fischer et al., 2022) as a multi-layer SpatRaster object at 10 × 10 km resolution in using the WGS84 coordinate reference system (EPSG:4326, “+proj=longlat +datum=WGS84 +no_defs”). Biome scheme numbers (1 to 31) follow Fischer et al. (2022). For each scheme, the metadata (original publication reference, spatial coverage, number and names of specific biomes, and scheme definition) are provided in the biomes_information() dataset and a summary can be obtained with the biomes_info() function (Table S1). Input occurrences can be provided as data.frame, sf or SpatVector object. If no occurrence data are available, biomes_occ() can take a taxon name, download occurrence records from GBIF and clean them, using *rgbif* and *CoordinateCleaner*. A vignette on how to obtain biome-data is provided on the webpage dedicated to the *biomes* package (https://azizka.github.io/biomes/articles/step1-assembly-of-occurrence-records-and-biome-schemes) and as additional .pdf for the peer review process.

#### Step 2: Choice of a biome scheme

biomes_rank() supports the selection of a suitable biome scheme by ranking the compared schemes according to their fit to a given occurrence dataset based on three data-driven criteria.

**1. Coverage** (*C*), the proportion of occurrence records that are located within any biome of the scheme.

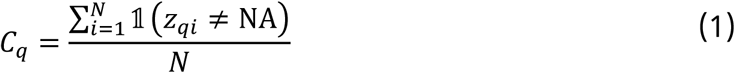

Here, *C_q_* is the Coverage of biome scheme *q; N* is the number of occurrence records with valid coordinates, and *i* indexes the individual records; *zqi* is the biome assigned to record *i* by scheme *q*, or NA if the record falls outside all biomes represented by that scheme. The indicator function equals 1 when the condition in parentheses is true and 0 otherwise. The numerator, denoted by *nq*, is therefore the total number of records assigned to any biome by scheme *q*.

**2. Effective number of biomes** (*E*), the Hill number of order 1, calculated as the exponential of the Shannon entropy of the distribution of assigned records among biomes:

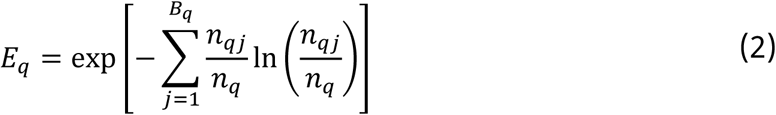

Here, *Eq* is the effective number of biomes for scheme *q*; *q* identifies the biome scheme; *Bq* is the total number of biomes in scheme *q*; and *j* indexes the individual biomes in scheme *q*; *nqj* is the number of records assigned to biome *j* in scheme *q*, whereas *nq* is the total number of records assigned to any biome in that scheme. Terms for which *nqj* = 0 contribute zero to the sum. *Eq* equals 1 when all assigned records occur in a single biome and increases as records are distributed more evenly across multiple biomes.

**3. Granularity** (*G*), is the proportion of the scheme’s biomes that contain at least one occurrence record:

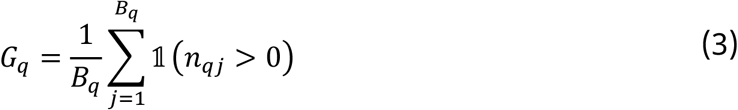

Here, *Gq* is the granularity of scheme *q*; *q* identifies the biome scheme; *Bq* is the total number of biomes in that scheme; and *j* indexes these biomes; *nqj* is the number of records assigned to biome *j* in scheme *q*. The indicator function 1(*nqj* > 0) equals 1 when biome *j* contains at least one occurrence record and 0 otherwise. Consequently, the sum in the numerator is the number of biomes in scheme *q* that contain at least one record, and division by *Bq* expresses this number as a proportion on the total number of biomes.

Each selected criterion X is rescaled linearly across the compared schemes:

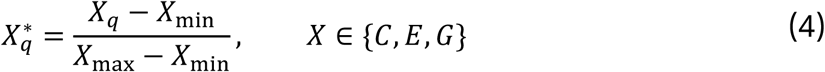

Here, *Xq* is the original value of criterion *X* for scheme *q*, *Xq\** is its rescaled value, and *Xmin* and *Xmax* are the minimum and maximum values of that criterion across the compared schemes. A rescaled value of 0 therefore represents the lowest value among the compared schemes rather than an absolute value of zero, while 1 represents the highest value.

The composite score is the unweighted mean of the rescaled criteria:

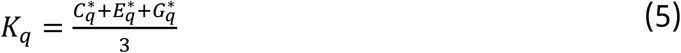

Here, *Kq* is the composite score of scheme *q*. If a criterion does not vary among the compared schemes, it is omitted from the numerator and the denominator is reduced accordingly. If none of the criteria varies, no composite score is defined.

The scheme with the highest *Kq* is returned as the top-ranked candidate. Ties in *Kq* are resolved by the publication year of the scheme (more recent schemes rank higher; tiebreaker argument). Because the rescaling in Equation 4 is relative to the compared set, *Kq* is comparable only among schemes included in the same ranking.

The ranking criteria provide a data-driven ranking that favours biome schemes that maximise coverage of the occurrence records while partitioning them among many evenly represented specific biomes as equally as possible. *E* and *G* measure different dimensions of the fit of a scheme to the data: *E* is bounded by the number of biomes a scheme distinguishes and therefore favours finely resolved schemes, whereas *G* favours coarsely resolved schemes in which few biomes remain empty. The composite score thus rewards the scheme whose resolution best matches the distribution of the records, rather than the finest or the coarsest scheme per se. However, it is still necessary for researchers to critically evaluate the suggestion in the light of their research question, and we recommend treating the resulting ranking as a shortlist. Limiting the comparison to a meaningful subset of schemes using the definition argument in the biomes_rank() function can be helpful to do so. The available subsets are “climate”, “vegetation”, “land_cover”, “ecoregion”, “integrative” and “anthropogenic”; setting definition = “all” (the default) ranks all 31 schemes. “Integrative” schemes synthesise multiple ecological or biogeographical criteria, concepts or data sources. Studies of climatic niche conservatism may prioritise climate-based schemes when climate is intended to define the biomes or exclude them when climate is the variable being tested, thereby avoiding circularity. Analyses constrained by classification coverage may instead select the conceptually appropriate scheme that classifies the largest proportion of records. Further information is provided in the “Choosing a biome scheme” vignette (https://azizka.github.io/biomes/articles/step2-choice-of-a-biome-scheme).

#### Step 3: Occurrences-to-biome classification

biomes_classify() uses the assembled data and selected biome scheme(s) to generate a data.frame with a biome assignment for each occurrence record. Classification can be run against any subset of schemes either using the scheme argument to select biome schemes by number (1 to 31), or the biome argument to pass a custom SpatRaster.

The function returns the occurrence data with one or two additional columns per scheme. Column names contain the raster-layer name followed by _name for the biome name or _value for its numerical raster code. The value argument determines which representation is returned. Occurrence records not covered by any biome in a given biome scheme, for instance on islands omitted from coarse-resolution maps, are assigned as “no_biome”. An alternative label can be specified using the na argument to the biomes_classify() function. Handling these records explicitly and identically across schemes matters, because the amount and spatial distribution of unassigned records differs among schemes and is one of the ranking criteria (Step 2). See the vignette “Occurrences-to-biome-classification” on GitHub (https://azizka.github.io/biomes/articles/step3-occurrence-to-biome-classification).

#### Step 4: Output and visualisation

biomes_tab() returns a table summarising the number of records assigned to each biome and biomes_visualise() creates a publication-style map showing the occurrence records over the selected scheme, with the option to include an inset summarising the proportion of records per biome. The resulting map is returned as a ggplot object, for customization with *ggplot2* syntax. See the vignette “Output and visualisation” for detailed examples of tabulating biome assignments and creating, customising, and exporting the resulting visualisations. (https://azizka.github.io/biomes/articles/step4-output-and-visualisation).

### Worked example: Bombacoideae

We demonstrate the workflow with 17,030 occurrence records from species in the plant subfamily Bombacoideae (Zizka et al., 2020), a group whose biome conservatism at the savanna-forest interface has been studied repeatedly (Pezzini et al., 2020; Zizka et al., 2020). Few lines of code are sufficient to run the full *biomes* workflow:

library(biomes)

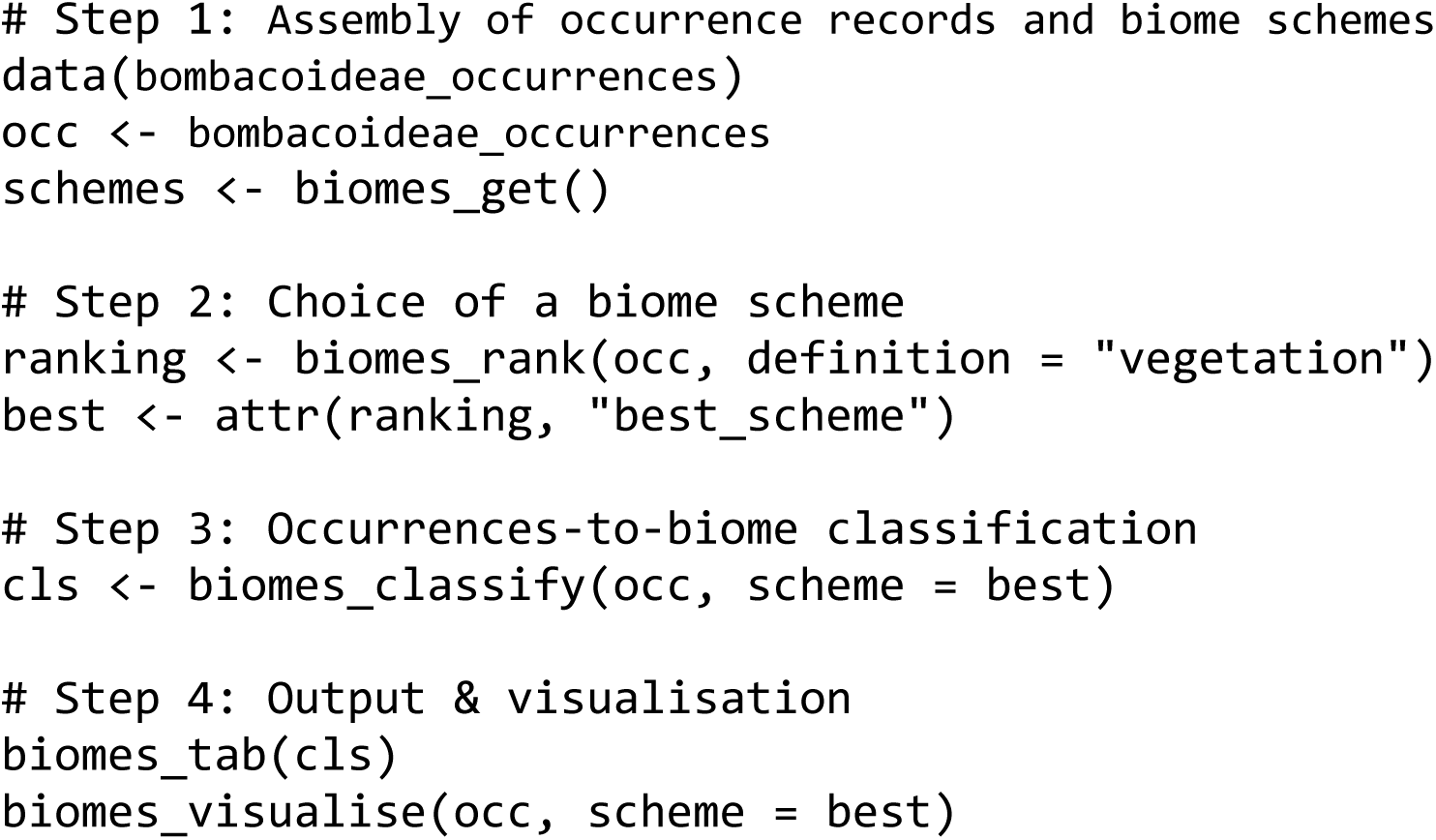

In the case of Bombacoideae, the relevant research question is about the adaptation to specific vegetation types (savanna vs. forest), thus a vegetation-based biome scheme is conceptually most adequate; we therefore rank within the “vegetation” biome definition (Table 1). An alternative unconstrained ranking across all 31 schemes is dominated by the anthropogenic-biome scheme of Ellis et al. (2010), because the near-complete coverage and many evenly occupied land-use classes maximise the data-driven criteria, but it answers a land-use rather than a vegetation adaptation question and is at odds with the age of many of the occurrence records, which pre-date the 1950ies. Restricting the ranking to the six vegetation-based schemes instead (Fig. 2a), selects the historical land-cover reconstruction of Ramankutty & Foley (1999), which approximates potential natural vegetation (Fig. 2b). This scheme ranks first for coverage and granularity, whereas its effective number of biomes is the lowest among the six schemes because it distinguishes only 12 biomes and about half of the records fall into a single biome (tropical evergreen woodland).

**Figure 2.**
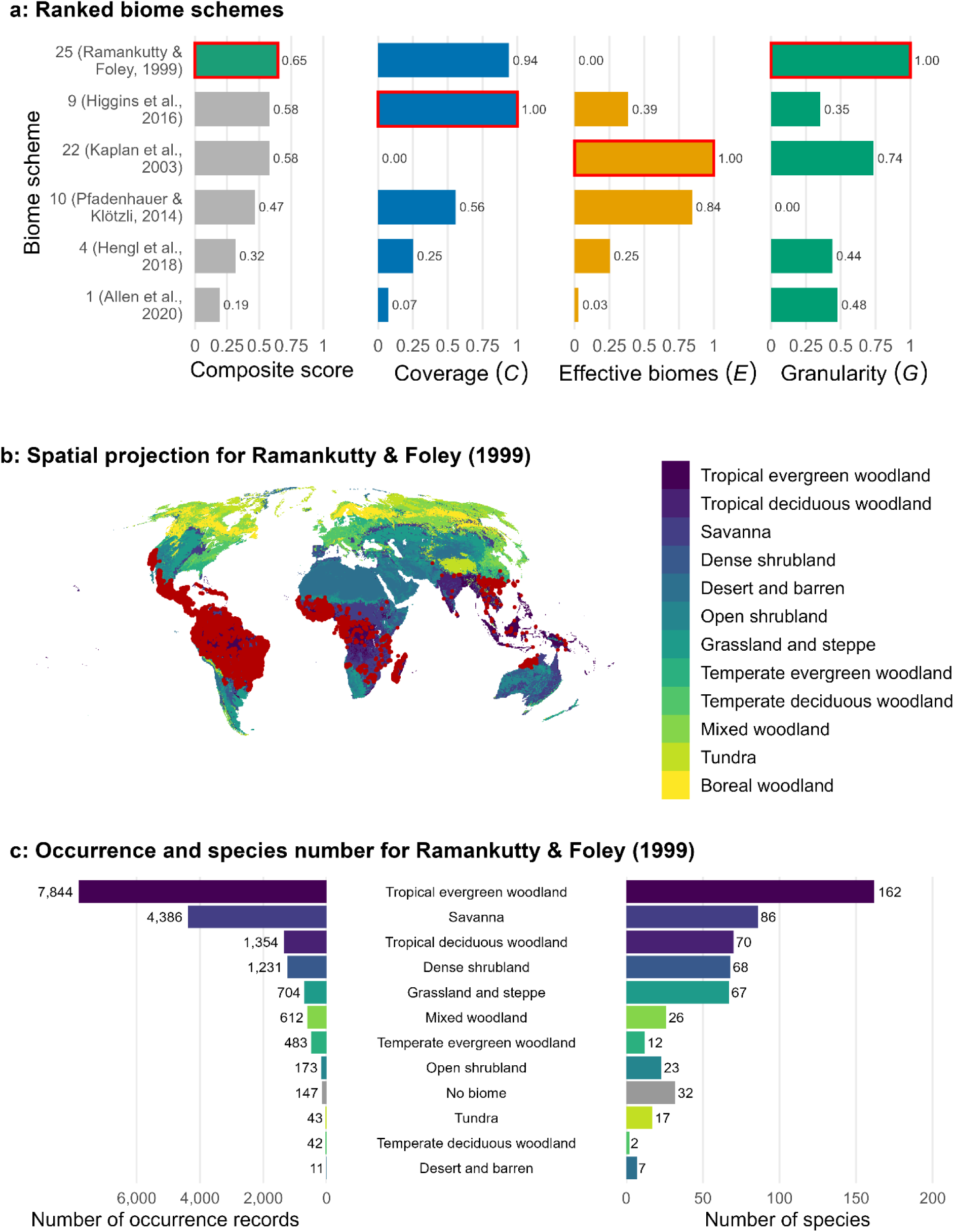
Standard visualization of the results from the *biomes* workflow, exemplified by a worked example of 17,030 occurrence records from 185 species in the Bombacoideae (Malavaceae). (a) Ranking of 6 vegetation-based biome schemes by suitability to the research question showing the composite score (*k*) per scheme as well as the individual scaled [0, 1] data-driven ranking criteria (*C, E, G*). Note that the scheme with the overall highest score has a low number of effective biomes, in this case a choice between the three best ranked biomes is reasonable. (b) Occurrence records mapped over the best-ranked scheme (Ramankutty & Foley, 1999). (c) The number of occurrence records (left) and species (right) per biome for the best-ranked scheme.

**Table 1.** The best-ranked biome scheme per biome definition, respectively for the Bombacoideae dataset from the worked example (biomes_rank()): the un-constrained ranking (“all”) points to an anthropogenic land-use scheme (no. 16), whereas the “vegetation” group selects biome scheme no. 25.

| Biome definition | No. biome schemes | Best biome Scheme source | Composite | % NA | Coverage | Effective biomes | Granularity |
| --- | --- | --- | --- | --- | --- | --- | --- |
| all | 31 | Ellis et al., 2010 | 0.97 | 1.12 | 0.99 | 12.42 | 1 |
| climate | 7 | Leemans, 1990 | 0.88 | 3.41 | 0.97 | 8.21 | 0.85 |
| vegetation | 6 | Ramankutty & Foley, 1999 | 0.65 | 0.86 | 0.99 | 4.65 | 0.92 |
| land_cover | 7 | Loveland et al., 2000 | 0.80 | 3.76 | 0.96 | 7.95 | 0.94 |
| ecoregion | 4 | The Nature Conservancy, 2009 | 0.70 | 3.01 | 0.97 | 4.61 | 0.88 |
| integrative | 6 | Schmithüsen, 1976 | 0.79 | 3.15 | 0.97 | 8.67 | 0.58 |
| anthropogenic | 1 | Ellis et al., 2010 | NA | 1.12 | 0.99 | 12.42 | 1 |

Under this scheme, Bombacoideae show the expected signal: “tropical evergreen woodland” accounts for 7,844 (46.1%) of the occurrence records and includes 162 of the 185 species (87.6%); “savanna” accounts for 4,386 (25.8%) of the records and includes 86 species (46.5%); and “tropical deciduous woodland” accounts for 1,354 (8.0%) of the records and includes 70 species (37.8%) (Fig. 2c). This is consistent with the biome-conservatism signal reported by Zizka et al. (2020), partly based on expert vegetation classification and shows that the standardised, definition-aware workflow recovers the expected biogeographic structure.

The *biomes* workflow facilitates the reproducible selection of biome schemes tailored to individual research questions. The top-ranked scheme depended on the type of biome definition considered (anthropogenic, climate, ecoregion, integrative, land cover, or vegetation), and within vegetation-based schemes the proportion of unassigned records ranged from 0.5% to 6.5% (Table 1; Fig. 2a). In our worked example, based on 17,030 occurrence records and restricted to the vegetation biome definition (definition = “vegetation”), Steps 1 to 3 were completed in approximately 3 s and Step 4 (tabulation and the three-panel figure) required an additional approximately 18 s on a personal laptop with an AMD Ryzen 5 5600H processor (6 cores, 12 threads) and 16 GB RAM.

## Conclusions

The *biomes* package addresses three challenges to the reproducible use of biomes in ecological and evolutionary research. First, it enables easy access to a compilation of 31 global biome schemes (Fischer et al., 2022) in a harmonised format within the R computational environment. Second, it facilitates a data-driven selection of the best-fitting biome scheme. Third, it implements a reproducible occurrence-to-biome classification with consistent handling of unassigned records. By lowering the technical barriers to working with multiple biome schemes, *biomes* can improve the efficiency, transparency and specificity of biome-based research in ecology, biogeography and evolutionary biology.

