## Supporting Information: Table S1 for "*biomes*: An R package for reproducible occurrence-to-biome classification using 31 global biome schemes"

**Table S1.** The 31 biome schemes provided by the *biomes* package. No.: raster-stack number; Name of biome scheme: scheme name; biome definition: the concept and data on which the biome scheme is based, one out of “anthropogenic”, “climate”, “ecoregion”, “integrative” (combined climate-vegetation schemes), “land_cover”, and “vegetation”; Literature reference: the reference to the original publication; No. of biomes: Total number of different biomes included in the scheme (zonal/azonal). Following Fischer et al. (2022), azonal biomes are not linked to climatic factors, such as inland water bodies, oceanic islands, mountains, and urban areas; Methodology (brief): a brief description of the applied methodology for biome delineation. See Fischer et al. (2022) for details.

| No. | Name of biome scheme | Biome definition | Literature reference | No. of biomes | Methodology (brief) |
|---|---|---|---|---|---|
| 1 | Global vegetation patterns of the past 140,000 years | vegetation | Allen et al., (2020) | 21 (21/0) | Modelling with the global dynamic vegetation Lund-Potsdam-Jena General Ecosystem Simulator. |
| 2 | Dataset of the global component of the Copernicus Land Monitoring Service | land_cover | Buchhorn et al., (2020) | 20 (18/2) | Supervised classification of satellite imagery data based on external reference data sets and expert opinion. |
| 3 | Present and future Köppen-Geiger climate classification maps at 1-km resolution | climate | Beck et al., (2018) | 30 (30/0) | Application of a slightly adjusted Köppen-Geiger classification based on climatological thresholds following Peel et al. (2007). |
| 4 | Global mapping of potential natural vegetation: an assessment of machine learning algorithms for estimating land potential | vegetation | Hengl et al., (2018) | 20 (20/0) | Modelling based on several machine learning techniques including neural networks, random forest, gradient boosting, K-nearest neighbour, and Cubist. |
| 5 | An ecoregion-based approach to protecting half the terrestrial realm | ecoregion | Dinerstein et al., (2017) | 14 (14/0) | Revision of the terrestrial ecoregions of the world by Olson et al. (2001) based on technical advances and expert knowledge. |
| 6 | A global classification of vegetation based on Normalized Difference Vegetation Index (NDVI), rainfall and temperature | integrative | Zhang et al., (2017) | 14 (14/0) | Non-hierarchical clustering based on K-means in 14 biomes, validation via Kappa statistics. |
| 7 | On using a clustering approach for global climate classification | climate | Netzel & Stepinski, (2016) | 13 (13/0) | Climate clustering based on dynamic time warping as a measure for dissimilarity between climate types. |
| 8 | On using a clustering approach for global climate classification | climate | Netzel & Stepinski, (2016) | 13 (13/0) | Climate clustering based on Euclidean distance as a measure |

| | | | | | |
|---|---|---|---|---|---|
| | | | | | for dissimilarity between climate types. |
| 9 | Defining functional biomes and monitoring their change globally | vegetation | Higgins et al., (2016) | 24 (24/0) | Vegetation categories based on multiple predefined parameters of vegetation height, productivity and plant growth limitations. |
| 10 | Earth´s vegetation | vegetation | Pfadenhauer & Klötzli, (2014) | 34 (31/3) | Review and modification of global vegetation patterns by Schmithüsen (1976) informed by multiple regional sources. |
| 11 | Spatiotemporal change in geographical distribution of global climate types in the context of climate warming | climate | Zhang & Yan, (2014) | 14 (14/0) | Non-hierarchical data clustering based on K-means distances in 14 classes, validation via Kappa statistics. |
| 12 | A high-resolution bioclimate map of the world: a unifying framework for global biodiversity research and monitoring | climate | Metzger et al., (2013) | 16 (16/0) | Compilation of multiple bioclimatic parameters, collinearity reduction among input parameters based on Pearson correlation, statistical grouping by principal components analysis of the covariance matrix, data clustering by iterative self-organizing data analysis for classification of principal components into homogeneous environmental strata, similarity-based aggregation of strata into global environmental zones based on Euclidean distance, comparison of final classification with multiple global and regional ecosystem concepts by Kappa statistics. |
| 13 | Global ecological zones for Food and Agriculture Organization (FAO) forest reporting: 2010 update | ecoregion | Food and Agriculture Organization of the United Nations, (2010) | 21 (20/1) | Global ecological zones based on a compilation of global and regional ecological and vegetational source maps, revision according to remote sensing. |
| 14 | Global Land Cover by national mapping organizations | land_cover | Tateishi et al., (2011, 2014); Kobayashi et al., (2017) | 19 (18/1) | Supervised classification of satellite imagery by Moderate Resolution Imaging Spectroradiometer (MODIS) based on multiple remote sensing products for reference as well as specific regional maps and expert opinion, individual unsupervised classification for certain classes, validation with stratified random sampling. |
| 15 | International Satellite Land Surface Climatology Project, Initiative II (ISLSCP II) University of Maryland global land cover classifications, 1992-1993 | land_cover | Defries et al., (2010) | 13 (12/1) | Resampling of land cover and derived Normalized Difference Vegetation Index (NDVI) data from Advanced Very High Resolution Radiometer (AVHRR) with hierarchical classification. |
| 16 | Anthropogenic transformation of biomes, 1700 to 2000 | anthropogenic | Ellis et al., (2010) | 19 (18/1) | Rule-based model classification according to standardized thresholds. |
| 17 | GlobCover | land_cover | European Space Agency, (2010) | 21 (20/1) | Regionally specified classification of high-resolution surface reflectance mosaics, validation informed by expert knowledge. |
| 18 | Moderate Resolution Imaging Spectroradiometer (MODIS) collection 5 global land cover: | land_cover | Friedl et al., (2010) | 16 (15/1) | Nested classification of Earth observation data based on ensemble decision trees, cross-validation analysis. |

| | | | | | |
|---|---|---|---|---|---|
| | algorithm refinements and characterization of new datasets | | | | |
| 19 | Terrestrial ecoregions of the world | ecoregion | The Nature Conservancy, (2009) | 16 (15/1) | Selected global and regional ecozones, alignment and expert informed modification. |
| 20 | Updated world map of the Köppen-Geiger climate classification | climate | Peel et al., (2007) | 31 (31/0) | Classification and spatial interpolation of observational climate records based on predefined thresholds. |
| 21 | Global Land Cover 2000 (GLC2000): a new approach to global land cover mapping from Earth observation data | land_cover | Bartholomé & Belward, (2005) | 21 (20/1) | Derivation of land cover maps from spectral surface reflectance at four wavelength ranges based on regionally optimized image classification procedure. |
| 22 | Climate change and Arctic ecosystems: 2. modeling, paleodata-model comparisons, and future projections | vegetation | Kaplan et al., (2003) | 27 (27/0) | Coupled biogeographic and biogeochemical distribution modelling of biomes of defined by main potential natural vegetation types. |
| 23 | Terrestrial ecoregions of the world: a new map of life on Earth | ecoregion | Olson et al., (2001) | 16 (15/1) | Review of global and regional biogeographic provinces, hierarchical classification into ecoregions, refinement based on expert consultation, nesting of ecoregions into biomes and biogeographic realms. |
| 24 | Development of a global land cover characteristics database and International Geosphere-Biosphere Programme (IGBP) Discover from 1 km Advanced Very High Resolution Radiometer (AVHRR) data | land_cover | Loveland et al., (2000) | 16 (15/1) | Unsupervised classification and subsequent stratification of monthly Normalized Difference Vegetation Index (NDVI) composites provided by AVHRR from 1992–1993 at continental scale. |
| 25 | Estimating historical changes in global land cover: croplands from 1700 to 1992 | vegetation | Ramankutty & Foley, (1999) | 12 (12/0) | Reclassification of 1-km DISCover land-cover data into potential natural vegetation types, aggregated to 5-arc-min cells using the dominant upland class and supplemented with modelled vegetation and climate-based rules in land-use-dominated areas. |
| 26 | Possible changes in natural vegetation patterns due to global warming | climate | Leemans, (1990) | 39 (38/1) | Application of the Holdridge life zone classification (Holdridge 1947, 1967) based on spatially interpolated bio-climatic parameters from climate station records. |
| 27 | Ecozones of the Earth | integrative | Schultz, (2016) | 15 (14/1) | Review, evaluation of regional ecological studies, quantitative ecosystem analysis. |
| 28 | Landscape belts of the Earth | integrative | Müller-Hohenstein, (1981) | 15 (13/2) | Review and combination of thematic concepts. |
| 29 | Atlas of biogeography | integrative | Schmithüsen, (1976) | 31 (29/2) | Biogeographical analysis integrating potential natural vegetation with climate (temperature and potential evapotranspiration), soils, topography, and elevation. |
| 30 | Communities and ecosystems | integrative | Whittaker, (1975) | 14 (12/2) | Graphical delineation of biome definition in climatic space defined by mean annual temperature and precipitation, |

| | | | | | |
|---|---|---|---|---|---|
| | | | | | informed by vegetation physiognomy and distribution. |
| 31 | Vegetation and climate | integrative | Walter, (1970); Breckle & Rafiqpoor, (2019) | 34 (32/2) | Classification according to defined thresholds on climatic data, vegetation proxies and surface cover indices. |
